# Variation in VKORC1 is associated with vascular dementia

**DOI:** 10.1101/2020.07.31.231126

**Authors:** Jure Mur, Daniel L. McCartney, Daniel I. Chasman, Peter M. Visscher, Graciela Muniz-Terrera, Simon R. Cox, Tom C. Russ, Riccardo E. Marioni

**Affiliations:** Lothian Birth Cohorts group, Department of Psychology, University of Edinburgh, Edinburgh, UK; Centre for Genomic and Experimental Medicine, Institute of Genetics and Molecular Medicine, University of Edinburgh, Edinburgh, UK; Alzheimer Scotland Dementia Research Centre, University of Edinburgh, Edinburgh, UK; Division of Preventive Medicine, Brigham and Women’s Hospital & Harvard Medical School, Boston, MA; Institute for Molecular Bioscience, University of Queensland, Brisbane, QLD, Australia; Edinburgh Dementia Prevention, University of Edinburgh, Edinburgh, UK; Division of Psychiatry, Centre for Clinical Brain Science, University of Edinburgh, UK

## Abstract

Genetic variation in *VKORC1* is associated with differences in the coagulation of blood and consequentially with sensitivity to the drug warfarin. Variation in *VKORC1* has also been linked to parental dementia. However, it is unclear whether the relationship persists for the diagnosis in patients themselves, whether the association holds only for certain forms of dementia, and if those taking warfarin are at greater risk. Here, we use data from 211,423 participants from UK Biobank to examine the relationship between *VKORC1*, risk of dementia, and the interplay with warfarin use. We find that the T-allele in rs9923231 confers a greater risk for vascular dementia (OR=1.28, p=0.0069), but not for general dementia (OR=1.04, p=0.21) or Alzheimer dementia (OR=1.05, p=0.35), and that the risk of vascular dementia is not affected by warfarin use in carriers of the T-allele. Our study reports for the first time an association between rs9923231 and vascular dementia, but further research is warranted to explore potential mechanisms and specify the relationship between rs9923231 and features of vascular dementia.

## Introduction

Warfarin is the most prescribed anticoagulant worldwide^1^ and is commonly used as a treatment for atrial fibrillation (AF)^2^. The drug functions by inhibiting the enzyme vitamin K epoxide reductase (VKOR), effectively interfering with the vitamin K cycle required for coagulation of blood^3^. As a result of variations in age, height, weight, genotype, and other factors^4–6^, patients vary up to 20-fold in their sensitivity to warfarin^7^. Clinically, the optimum dose is estimated using tests of blood coagulation, commonly the International Normalised Ratio (INR). The strongest genetic predictor of warfarin sensitivity is the gene *VKORC1*, which encodes for the vitamin K epoxide reductase subunit 1 (VKORC1) and accounts for approximately a third of the variance in warfarin sensitivity^3^. Three *VKORC1* SNPs, rs9923231, rs9934438, and rs2359612 – which are in very high linkage disequilibrium – are the best genetic predictors of warfarin sensitivity^3,7,8^.

In a recent GWAS meta-analysis of parental dementia and case-control Alzheimer dementia (ADem)^9^, *VKORC1* was associated (after Bonferroni correction) with ADem in a gene-based test (p=5.1×10^−8^); the T-allele in rs9923231 – which is related to the need for a lower dose of warfarin – was not a genome-wide significant finding, but was both located within a genome-wide significant locus and nominally associated with an increased risk of ADem (p=1.8×10^−7^). Pure Alzheimer’s disease pathology – characterised by amyloid plaques and neurofibrillary tangles in the grey matter – is uncommon, and most patients exhibit a mixed pathology in which vascular factors often play a prominent role^10^. In fact, there is extensive evidence directly linking vascular dysfunction to ADem^11^. Thus, a possible explanation for the findings^9^ is that vascular factors played a crucial role in a proportion of the ADem cases/family history cases observed. If that is the case, then there should be an even stronger relationship between *VKORC1* and vascular dementia (VaD) that is mostly due to cardiovascular factors.

Most strokes in western countries are due to occlusions in blood vessels (ischaemic), and some are due to ruptures in blood vessels (haemorrhagic)^12^. If carriers of the T-allele in rs9923231 experience a reduction of blood coagulation and subsequent sequential minor haemorrhagic strokes, the resulting pathology could manifest in dementia and explain the observed link. Furthermore, compared to non-carriers of the T-allele, patients with AF that carry the T-allele could be at an increased risk of intracerebral haemorrhage and consequentially vascular dementia (VaD) when prescribed warfarin. Here, we study the same UK Biobank cohort as previously^9^, but consider both individual and parental dementia status. We test whether T-allele status is associated with an increased risk of VaD and explore whether carriers of the T-allele are at a greater risk of VaD than non-carriers when prescribed warfarin.

## Methods

### Sample

We used data from UK Biobank, a large and detailed prospective study of over 500,000 participants aged 37-73 that were recruited between the years 2006 and 2010. UK Biobank has been described in detail before^13^. The Research Ethics Committee (REC) granted ethical approval for the study— reference 11/NW/0382—and the current analysis was conducted under data application 10279.

### Genotyping

Details on genotyping in the UK Biobank have been reported before^14,15^. Briefly, for 49,950 participants, genotyping was performed using the UK BiLEVE Axiom Array, and for 438,427 participants, genotyping was performed using the UK Biobank Axiom Array. The released data contained 805,426 markers for 488,377 participants. Further quality control steps were performed as previously reported^9^. They included the removal of outliers, of incongruent data points, and of related participants using a relationship cut-off of 0.025 (GCTA GREML)^16^. This left an unrelated cohort of 332,050 individuals of white British ancestries (**Figure 1**).

**Figure 1:**
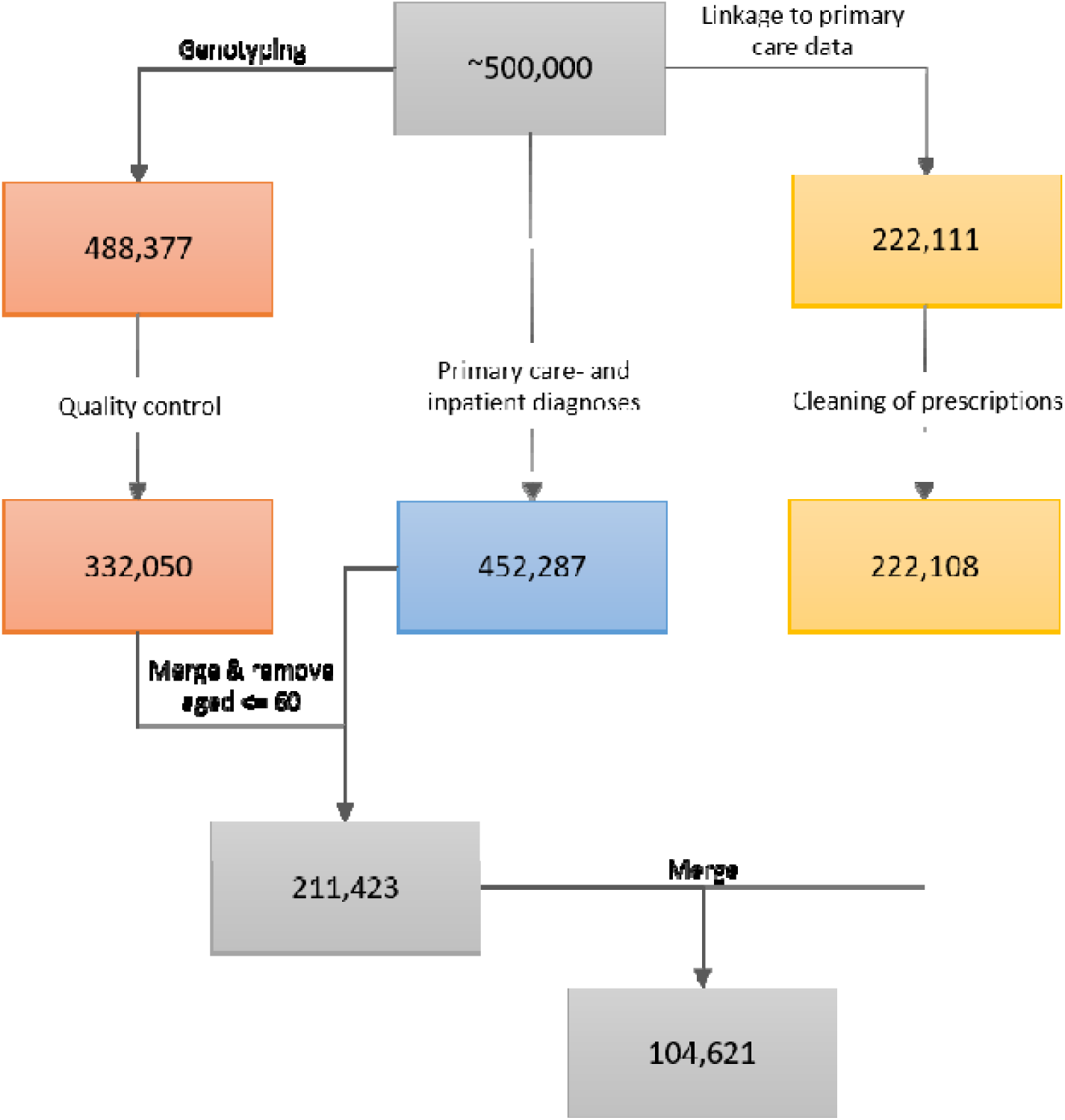
The data cleaning procedure. The left path (orange boxes) represents the genotyping and associated quality control; the middle path (blue box) represents the ascertainment of primary care- and inpatient diagnoses; the right path (yellow boxes) represents the linkage to primary care prescriptions and the cleaning of the latter. The last two steps (grey boxes) involved the inclusion of only those participants that were older than 60 at the end of sampling and who passed through the left and middle paths (first grey box, 211,423 participants), or through all three data-cleaning paths (second grey box, 104,621 participants). All analyses that did not include prescribing data in the models, were performed using the 211,423 participants, while the analyses that utilized warfarin

### Warfarin prescription data

The UK Biobank obtained data on prescriptions for 222,111 participants via primary care computer system suppliers (EMIS Health and Vision for Scotland, and Wales, Vision and The Phoenix Partnership for England) and has engaged other intermediaries (Albasoft – a third party data processor – for Scotland and the SAIL databank for Wales). All participants provided written consent for linkage to their health records upon recruitment to UK Biobank. The data were extracted in May 2017 for Scotland, in September 2017 for Wales, and in June, in July, and in August 2017 for England. The data include the exact dates of prescriptions, drug codes (BNF, Read v2, CTV3, and dm+d), names of drugs as written on the prescription, and – where available – the dosages of prescribed drugs. Empty prescriptions, prescriptions without a date, and duplicate prescriptions (defined as identical prescriptions issued to the same person on the same day) were removed from the sample. This resulted in the removal of 1,467,547 prescriptions. Three participants were completely removed from the dataset (Figure 1). Warfarin prescriptions were extracted by searching for the word “warfarin” under the name/content of each prescription. We calculated warfarin use for each participant by summing the number of days on which warfarin was prescribed.

### Disease status

Data on diagnoses for 452,287 participants were obtained by the UK Biobank from two sources: (1) from primary care similarly to the prescriptions described above, and (2) from hospital inpatient admissions data. Inpatients are defined as people who are admitted to hospital and occupy a hospital bed. These data included Hospital Episode Statistics for England, Scottish Morbidity Records for Scotland, and the Patient Episode Database for Wales. People with record of any dementia were included in a broad dementia category of “general dementia”, and narrower, more specific categories-ADem, VaD – were also identified. Information on the codes used in the extraction of diagnoses is provided in **Suppl. Table 1**. We excluded from our analyses all participants that were 60 years old or younger on the last date of sampling (27.09.2017) since dementia risk increases steeply with age. Parental diagnoses were ascertained during the initial assessment by asking participants about the presence of “Alzheimer’s disease/dementia” for both mother and father. In our analyses, the parental diagnosis of dementia was considered positive if at least one parent was reported to have suffered from the disorder.

**Table 1:** Demographic characteristics of the sample.

| Variable | Level | Median (IQR) or n (%) |  |  |  |
| --- | --- | --- | --- | --- | --- |
|  |  | All<br>(n=211,423) | General dementia<br>(n=2,407) | ADem<br>(844) | VaD<br>(301) |
| Age |  | 61.0 (7.9) | 65.3 (6.1) | 65.6 (5.7) | 66.6 (4.7) |
| Sex | Female | 114,036 (53.9) | 1,054 (43.8) | 438 (51.9) | 110 (36.5) |
|  | Male | 97,387 (46.1) | 1,353 (56.2) | 406 (48.1) | 191 (63.5) |
| Education | Graduate degree | 62,670 (29.9) | 553 (23.4) | 170 (20.6) | 50 (17.1) |
|  | No graduate degree | 146,653 (70.1) | 1,808 (76.6) | 656 (79.4) | 242 (82.9) |
| Deprivation |  | -2.5 (3.6) | -2.3 (4.2) | -2.3 (4.0) | -1.9 (4.4) |
| Alcohol consumption | Daily or almost daily | 48,776 (23.1) | 549 (22.8) | 165 (19.5) | 67 (22.3) |
|  | 3 or 4 times a week | 50,390 (23.8) | 474 (19.7) | 173 (20.5) | 53 (17.6) |
|  | 1 or 2 times a week | 52,685 (24.9) | 535 (22.2) | 192 (22.7) | 74 (24.6) |
|  | 1-3 times a month | 21,266 (10.3) | 216 (9.0) | 83 (9.8) | 19 (6.3) |
|  | Special occasions only | 22,891 (10.8) | 309 (12.8) | 116 (13.7) | 41 (13.6) |
|  | Never | 14,882 (7.0) | 322 (13.4) | 115 (13.6) | 47 (15.6) |
| Smoking | Current smoker | 18,589 (8.8) | 223 (9.3) | 58 (6.9) | 39 (13.0) |
|  | Previous smoker | 81,754 (38.8) | 1,016 (42.5) | 351 (41.9) | 142 (47.3) |
|  | Non-smoker | 110,305 (52.4) | 1,154 (48.2) | 429 (51.2) | 119 (39.7) |
| Physical activity | Strenuous | 15,684 (7.9) | 106 (4.9) | 45 (5.8) | 4 (1.6) |
|  | Moderate | 132,566 (66.9) | 1,314 (61.1) | 482 (62.0) | 157 (60.9) |
|  | Light | 49,880 (25.2) | 730 (34.0) | 251 (32.3) | 97 (37.6) |
| BMI |  | 27.0 (5.7) | 27.1 (5.8) | 27.0 (5.6) | 27.5 (7.1) |

### Models

All analyses where the outcome variable was continuous were performed using linear regression; all models where the outcome variable was binary were performed using logistic regression. All models were controlled for the assessment centre in which the participant was tested, the genotyping-batch and array, 40 genetic principal components, the age, sex, education, deprivation, alcohol consumption, smoking, physical activity, and BMI of the participants. All covariates were ascertained immediately prior to or during the participants’ recruitment to the UK Biobank. For education, a binary classification was used that indicated whether a graduate degree had been attained. For socioeconomic deprivation, the Townsend index^17^ was used, where higher values indicate greater deprivation (range in the sample: −6.3-10.8). For alcohol consumption, a 6-level scale of frequency of alcohol consumption was used, where 1: “daily or almost daily”, 2: “three or four times a week”, 3: “one or two times a week”, 4: “one to three times a month”, 5: “special occasions only”, 6: “never”. For smoking, the participants were classified as non-smokers, past smokers, or current smokers. For physical activity, the scale provided by the UK Biobank was reduced to a 3-level scale, indicating light, moderate, or strenuous physical activity, as has been used before^18^. For analyses where parental diagnoses were modelled as outcomes, the ages of each parent (current age or age at death) were included in the models. In all cases where we tested for associations between rs9923231 (*VKORC1*) and any form of dementia, we assumed an additive genetic effect for rs99232331. All covariates were simultaneously added to the model and the models were not corrected for multiple comparisons. All analyses were performed in R version 3.6.3. The code for preparing and analysing the data is available at https://github.com/Logos24/VKORC1-and-VaD.

## Results

### Sample characteristics

Among the 211,423 participants, 114,036 (53.9%) were female and 97,387 (46.1%) were male (Table 1). The age range at recruitment was 48-73 years (Figure 2) and the median age was 61.0 years (IQR=7.9). A total of 4,059 (1.9%) participants had been diagnosed with AF and among the 104,621 participants with data on prescriptions, 5,331 (5.1%) had a history of being prescribed warfarin (**Suppl. Table 2**).

**Table 2:** Results of the additive models with T as the effect allele, using rs9923231 to predict parental dementia, general dementia, ADem, and VaD.

| Effect per T allele |  |  |  |  |
| --- | --- | --- | --- | --- |
| rs9923231 | OR | 95% CI | p | n cases |
| Parental dementia | 1.04 | 1.02-1.06 | $2.1 \times 10^{-4}$ | 30,626 |
| General dementia | 1.04 | 0.98-1.11 | 0.21 | 2,407 |
| ADem | 1.05 | 0.95-1.17 | 0.35 | 844 |
| VaD | 1.28 | 1.07-1.53 | 0.0069 | 301 |

**Figure 2:**
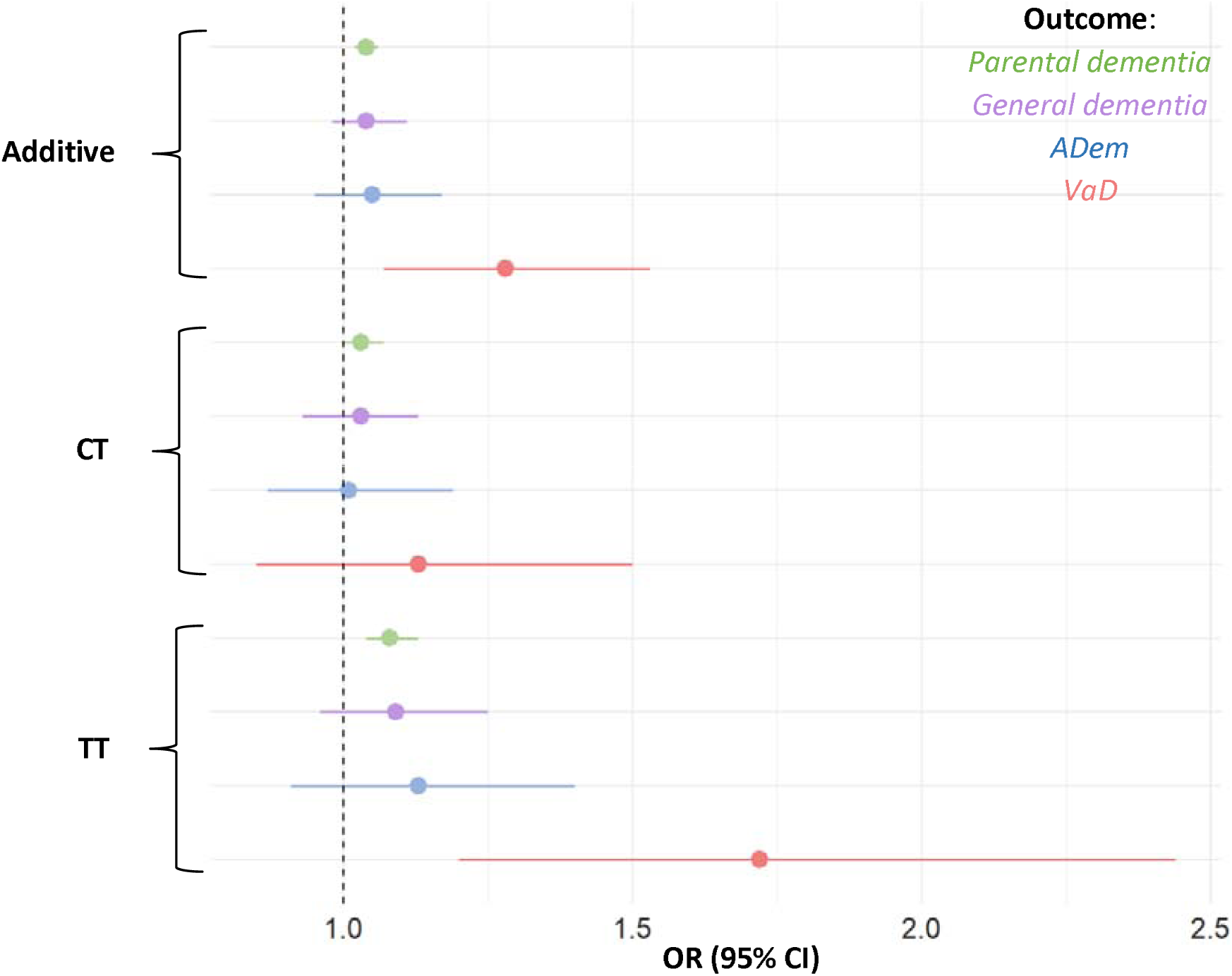
Odds ratios for parental dementia, ADem, and VaD per rs9923231 genotype status. Depicted are the additive effect and the effects of each allele group. The tails represent 95% confidence intervals for the OR’s.

There were 128,711 (60.9%) carriers of the T-allele in the sample: 99,117 (46.9%) were heterozygous for the T-allele, and 29,594 (14.0%) were homozygous for the T-allele; the allele frequencies were in Hardy-Weinberg equilibrium (*χ*^2^ =0.13, df=1, p=0.72). Among the participants, 2,407 (1.1%) had been diagnosed with general dementia, 844 (0.40%) with ADem (**Suppl. Table 3, Suppl. Figure 1**), and 301 0.14%) with VaD (**Suppl. Table 3, Suppl. Figure 1**); 58 participants (0.03%) had been diagnosed with both ADem and VaD. Patients with at least one parent with dementia were more likely develop ADem (OR=3.6, 95% CI=3.0-4.4, p<2.0×10^−16^) and more likely to develop VaD (OR=3.0, 95% CI=2.1- 4.3, p<1.2×10^−9^).

### rs9923231 polymorphism and warfarin dose

Carrying the T-allele was negatively associated with the average dose of warfarin (beta_per T-allele_=−0.29, SE=0.020, p<2.0×10^−16^). Individuals heterozygous for the T-allele were prescribed a dose of warfarin that was on average 0.23mg smaller than the dose prescribed to non-carriers (SE=0.022, p<2.0×10^−16^), while individuals homozygous for the T-allele were prescribed a dose of warfarin that was on average 0.62mg smaller than the dose prescribed to non-carriers (SE=0.033, p<2.0×10^−16^). The average dose of warfarin was also negatively associated with age (beta=−0.011, SE=0.0023, p=2.6×10^−6^), and was higher in males (beta=0.058, SE=0.023, p=0.011).

### rs9923231 polymorphism and dementia risk

Parents of carriers of the T-allele were more likely to have developed dementia (additive effect per T-allele: OR=1.04, 95% CI=1.02-1.06, p=2.1×10^−4^). When the presence of the T-allele was used to predict general dementia in participants, the effect was not significant, nor was the effect significant when the presence of the T-allele was used to predict ADem in participants **(Table 2, Figure 2)**.

When limited to the specific outcome of VaD, the additive effect of the T-allele was significant (OR=1.28, 95% CI=1.07-1.53, p=0.0069, Table 2, Figure 2). The full breakdown of all allele groups is shown in **Suppl. Table 4**. The sizes of the effects were numerically larger than for the association between the *VKORC1* genotype and either parental dementia, general dementia in participants, or ADem in participants. We repeated the models for VaD, with rs9923231 as a predictor and with the simultaneous addition of diagnoses of hypertension (n=69,005) and hypercholesterolemia (n=31,174) as additional covariates. While both hypertension and hypercholesterolemia were significant predictors (p=4.8×10^−14^ and p=3.8×10^−5^, respectively), this did not affect the relationship between rs9923231 and VaD (**Suppl. Table 5**). *Warfarin use and VaD in carriers of the T-allele*

In our sample, participants diagnosed with AF were not at greater risk for ADem compared to those without AF (OR=0.95, 95% CI=0.60-1.41, p=0.81). Participants diagnosed with AF were at a greater risk for VaD compared to those without AF (OR=2.32, 95% CI=1.35-3.76, p=1.1×10^−3^). The effect remained significant when hypercholesterolemia and hypertension were included in the model as covariates (OR=1.80, 95% CI=1.04-2.90, p=0.024). To test whether warfarin use in T-allele carriers diagnosed with AF increases the risk of VaD, we performed a logistic model with AF, warfarin use, and rs9923231 predicting VaD, with the inclusion of a 3-way interaction term between AF, warfarin, and rs9923231. The interaction between AF, warfarin use, and rs9923231 was not significant (OR=1.0, 95 % CI=0.97-1.01, p=0.38). The two-way interactions between the above variables were also not significant and effect sizes (main effects) were not substantially attenuated by the addition of the other variables into the models (**Suppl. Table 6, Suppl. Table 7**). Due to the small number of people with VaD and very limited statistical power for these analyses (**Suppl. Text 1**), we repeated the analysis by modelling parental dementia as an outcome and including the 3-way interaction term as above; parental dementia was thus treated as a proxy for VaD in the participants. The interaction between AF, warfarin use, and carrier-status was not significant (OR=1.0, 95% CI=0.996-1.01, p=0.79).

## Discussion

In this study, we explored the relationship between dementia, atrial fibrillation, warfarin use, and rs9923231, whose T-allele is associated with a reduction in the dose of warfarin^3,7,8^. We found a significant association between rs9923231 and VaD, but not between rs9923231 and either general dementia or ADem. While AF was linked to AF, the use of warfarin in patients that have AF and carry the T-allele did not increase the risk for VaD.

While there have been reports of variants for monogenic forms of VaD^19^, data on the genetics of sporadic VaD are sparse. To our knowledge, only two GWAS’s have been conducted to investigate this: One (n=5,700)^20^ found only rs12007229 on the X-chromosome to be linked to incident VaD, while the other (n=284)^21^ did not find any significant associations for VaD. A systematic review of all genetic association studies for the broader term of vascular cognitive impairment (VCI) found an association for 6 SNPs in 6 genes: *APOE, ACT, ACE, MTHFR, PON1*, and *PSEN-1*^*22*^.

Previous research has associated variation in rs9923231 with warfarin dose^3,7,8,23^, and with various adiposity-related traits, such as hip circumference, arm- and leg fat mass, and BMI (Gene Atlas^24^). To our knowledge the present study for the first time describes an association between rs9923231 and VaD, although it is important to note that this is not at a genome-wide significant threshold. The lack of a relationship between rs9923231 and either ADem or general dementia in the present study suggests that the association between the T-allele and ADem – as reported previously^9^ – might have been partly due to the classification of parental dementia. The UK Biobank questionnaire administered to participants did not distinguish between different types of dementia and it is not known how many of the 42,034 parents that were reportedly diagnoses with “Alzheimer’s/dementia”^9^ may have suffered from VaD. This hypothesis is further supported by the estimated effect sizes for the association between rs9923231 and ADem, which were not numerically larger than those for the association between rs9923231 genotype and parental dementia. Since parental dementia was used as a proxy for ADem in participants, the effect for ADem in participants should have been substantially greater than for parental dementia (even in the absence of statistical significance) if there truly was an association between rs9923231 and ADem (as opposed to an association between rs9923231 and VaD). Furthermore, in a recent GWAS of clinically diagnosed ADem (n=94,437)^25^ there was no association between rs9923231 and ADem.

Based on our results and considering the importance of cardiovascular abnormalities in the pathology of dementia^10,11^, any future studies exploring the association between rs9923231 and dementia must strongly consider the role of cardiovascular factors: The relationship between genotype and dementia might hold only for cases of pure VaD or for those in which vascular pathology represents the main cause of the disorder.

### Interplay between AF, VaD and warfarin use

AF has been previously associated with cognitive decline and dementia. Despite a substantial overlap of risk factors for AF and ADem, there is some evidence for an independent relationship between the two disorders^26,27^. In our study, AF was associated with VaD, even after controlling for hypertension and hypercholesterolemia. The association between AF and VaD is unsurprising, considering the inclusion of either vascular disease or history of stroke in almost all definitions of VaD^28^. However, we did not find an association between AF and ADem.

Due to the positive association between AF and VaD, the relationship between rs9923231 and VaD, and between rs9923231 and required warfarin dose, T-allele carriers that take warfarin to treat their AF might be at an increased risk of VaD than non-carriers due to warfarin-related brain haemorrhages. To test this, we studied an interaction between warfarin use, AF, and *VKORC1* genotype with VaD. We observed no variation in dementia risk by different combinations of these predictors. Due to reduced coagulation of blood in carriers of the T-allele, these individuals could be at greater risk of internal bleeding when taking warfarin. However, the required dose of warfarin is regularly estimated and adjusted using tests of blood coagulation and based on the results of the present paper, this approach is just as efficient in patients carrying the T-allele.

### Limitations and future directions

The present study has the advantages of having used a well-characterised sample with access to both inpatient- and primary-care diagnoses. However, we acknowledge several limitations. Despite the large number of people recruited to UK Biobank, the age range at the end of the sampling period for the cohort is 60-83 years, resulting in a low incidence and prevalence of dementia. This heavily reduced the size of our sample, especially when testing for interactions, and led to wide confidence intervals for the estimated odds ratios. Furthermore, the dose of warfarin ingested by participants was assumed to correspond to the average of their prescribed dose, despite some individuals possibly taking more or less of the medicine depending on their individual drug regimes. Finally, while most definitions of VaD include both dementia and a history of stroke or cardiovascular disease, VaD is very heterogeneous^29^; in the present study, we did not explore potential mechanisms and mediators of the association between rs9923231 and VaD, nor did we test the relationship for different subtypes of VaD.

The knowledge of genetic risk factors for diseases enables the generation of more accurate hypotheses about underlying biological mechanisms and illuminates potential targets for pharmacological intervention. Moreover, it allows for more informed stratification of participants in clinical trials. Studies that build on our research should aim to replicate the findings in a bigger sample and with greater precision determine the effect size for the association between rs9923231 and VaD. Additionally, further work is required to identify possible associations between rs9923231 and features of VaD, such as lacunar infarction, intracerebral haemorrhage, and white matter hyperintensities.

## Acknowledgements

DLM and REM are supported by Alzheimer’s Research UK major project grant ARUK-PG2017B-10. JM is supported by funding from the Wellcome Trust 4-year PhD in Translational Neuroscience—training the next generation of basic neuroscientists to embrace clinical research [108890/Z/15/Z]. PMV acknowledges funding from the Australian National Health and Medical Research Council (1113400) and the Australian Research Council (FL180100072). JM and TCR are members of the Alzheimer Scotland Dementia Research Centre funded by Alzheimer Scotland. TCR is employed by NHS Lothian and the Scottish Government. SRC is supported by Age UK (Disconnected Mind project), the UK Medical Research Council [MR/R024065/1] and a National Institutes of Health (NIH) research grant R01AG054628. The authors thank Dr Michelle Luciano (Department of Psychology, University of Edinburgh) for managing UK Biobank data application 10279.

## References

1. Ross, K. A. et al. Worldwide allele frequency distribution of four polymorphisms associated with warfarin dose requirements. J. Hum. Genet. 55, 582–589 (2010).

2. Zimetbaum, P. Atrial Fibrillation. Ann. Intern. Med. 166, ITC33–ITC48 (2017).

3. Wadelius, M. et al. Common VKORC1 and GGCX polymorphisms associated with warfarin dose. Pharmacogenomics J. 5, 262–270 (2005).

4. Saleh, M. I. Clinical predictors associated with warfarin sensitivity. Am. J. Ther. 23, e1690–e1694 (2016).

5. Sconce, E. A. et al. The impact of CYP2C9 and VKORC1 genetic polymorphism and patient characteristics upon warfarin dose requirements: proposal for a new dosing regimen. Hemostasis, Thromb. Vasc. Biol. 106, 2329–2333 (2005).

6. Wells, P. S., Holbrook, A. M., Crowther, N. R. & Hirsh, J. Interactions of warfarin with drugs and food. Ann. Intern. Med. 121, 676–683 (1994).

7. Takeuchi, F. et al. A genome-wide association study confirms VKORC1, CYP2C9, and CYP4F2 as principal genetic determinants of warfarin dose. PLoS Genet. 5, (2009).

8. Cha, P. C. et al. Genome-wide association study identifies genetic determinants of warfarin responsiveness for Japanese. Hum. Mol. Genet. 19, 4735–4744 (2010).

9. Marioni, R. E. et al. GWAS on family history of Alzheimer’s disease. Transl. Psychiatry 8, 0–6 (2018).

10. Van Der Flier, W. M. et al. Vascular cognitive impairment. Nat. Rev. Dis. Prim. 4, 1–16 (2018).

11. Sweeney, M. D. et al. Vascular dysfunction—The disregarded partner of Alzheimer’s disease. Alzheimer’s Dement. 15, 158–167 (2019).

12. Kalaria, R. N., Akinyemi, R. & Ihara, M. Stroke injury, cognitive impairment and vascular dementia. Biochim. Biophys. Acta – Mol. Basis Dis. 1862, 915–925 (2016).

13. Sudlow, C. et al. UK Biobank: An Open Access Resource for Identifying the Causes of a Wide Range of Complex Diseases of Middle and Old Age. PLoS Med. 12, 1–10 (2015).

14. Bycroft, C. et al. The UK Biobank resource with deep phenotyping and genomic data. Nature 562, 203–209 (2018).

15. Wain, L. V. et al. Novel insights into the genetics of smoking behaviour, lung function, and chronic obstructive pulmonary disease (UK BiLEVE): A genetic association study in UK Biobank. Lancet Respir. Med. 3, 769–781 (2015).

16. Yang, J., Lee, S. H., Goddard, M. E. & Visscher, P. M. GCTA: A tool for genome-wide complex trait analysis. Am. J. Hum. Genet. 88, 76–82 (2011).

17. Townsend, P. Deprivation. J. Soc. Policy 16, 125–146 (1987).

18. Quinn, T. J., Gallacher, K. I., Neal, S. R., Lee, D. & Mair, F. S. Assessing Risks of Polypharmacy Involving Medications With Anticholinergic Properties. Ann. Fam. Med. 18, 148–155 (2020).

19. Ikram, M. A. et al. Genetics of vascular dementia – review from the ICVD working group. BMC Med. 15, 1–7 (2017).

20. Schrijvers, E. M. C. et al. Genome-wide association study of vascular dementia. Stroke 43, 315–319 (2012).

21. Kim, Y., Kong, M. & Lee, C. Association of intronic sequence variant in the gene encoding spleen tyronase kinase with susceptability to vascular dementia. World J. Biol. Psychiatry 14, 220–226 (2013).

22. Dwyer, R., Skrobot, O. A., Dwyer, J., Munafo, M. & Kehoe, P. G. Using Alzgene-like approaches to investigate susceptibility genes for vascular cognitive impairment. J. Alzheimer’s Dis. 34, 145–154 (2013).

23. Buniello, A. et al. The NHGRI-EBI GWAS Catalog of published genome-wide association studies, targeted arrays and summary statistics 2019. Nucleic Acids Res. 47, D1005–D1012 (2019).

24. Canela-Xandri, O., Rawlik, K. & Tenesa, A. An atlas of genetic associations in UK Biobank. Nat. Genet. 50, 1593–1599 (2018).

25. Kunkle, B. W. et al. Genetic meta-analysis of diagnosed Alzheimer’s disease identifies new risk loci and implicates Aβ, tau, immunity and lipid processing. Nat. Genet. 51, 414–430 (2019).

26. Aldrugh, S., Sardana, M., Henninger, N., Saczynski, J. S. & McManus, D. D. Atrial fibrillation, cognition and dementia: A review. J. Cardiovasc. Electrophysiol. 28, 958–965 (2017).

27. Dietzel, J., Haeusler, K. G. & Endres, M. Does atrial fibrillation cause cognitive decline and dementia? Europace 20, 408–419 (2018).

28. Gorelick, P. B. et al. Vascular contributions to cognitive impairment and dementia: A statement for healthcare professionals from the American Heart Association/American Stroke Association. Stroke 42, 2672–2713 (2011).

29. Perneczky, R. et al. Is the time ripe for new diagnostic criteria of cognitive impairment due to cerebrovascular disease? Consensus report of the International Congress on Vascular Dementia working group. BMC Med. 14, 1–10 (2016).

