## Supplementary material for "Variation in VKORC1 is associated with vascular dementia": Suppl. material

| **Disorder** | **Code** | **Code system** | **Source** | **n** |
| --- | --- | --- | --- | --- |
| ADem | XaIkh | Read code | GP | 184 |
| ADem | Eu00. | Read code | GP | 143 |
| ADem | Eu002 | Read code | GP | 46 |
| ADem | Eu00z | Read code | GP | 16 |
| ADem | Fyu30 | Read code | GP | 1 |
| ADem | F110. | Read code | GP | 283 |
| ADem | F1101 | Read code | GP | 1 |
| ADem | X002x | Read code | GP | 32 |
| ADem | XaIKB | Read code | GP | 24 |
| ADem | X0030 | Read code | GP | 37 |
| ADem | X00R2 | Read code | GP | 25 |
| ADem | XaIKC | Read code | GP | 6 |
| ADem | Y016b | Read code | GP | 65 |
| ADem | Y016d | Read code | GP | 3 |
| ADem | E001. | Read code | GP | 6 |
| ADem | E001z | Read code | GP | 3 |
| ADem | E000. | Read code | GP | 1 |
| ADem | E0020 | Read code | GP | 1 |
| ADem | G309 | ICD10 | Inpatient | 815 |
| ADem | F009 | ICD10 | Inpatient | 457 |
| ADem | F002 | ICD10 | Inpatient | 94 |
| ADem | G308 | ICD10 | Inpatient | 90 |
| ADem | G301 | ICD10 | Inpatient | 41 |
| ADem | F001 | ICD10 | Inpatient | 37 |
| ADem | G300 | ICD10 | Inpatient | 95 |
| ADem | F000 | ICD10 | Inpatient | 60 |
| ADem | 2901 | ICD9 | Inpatient | 1 |
| AF | G573z | Read code | GP | 51 |
| AF | G5732 | Read code | GP | 1 |
| AF | Xa2E8 | Read code | GP | 1773 |
| AF | XaOfa | Read code | GP | 59 |
| AF | XaOft | Read code | GP | 33 |
| AF | X202S | Read code | GP | 3 |
| AF | XaeUP | Read code | GP | 1 |
| AF | G5730 | Read code | GP | 4160 |
| AF | G573. | Read code | GP | 523 |
| AF | I480 | ICD10 | Inpatient | 1306 |
| AF | I481 | ICD10 | Inpatient | 215 |
| AF | I482 | ICD10 | Inpatient | 68 |
| AF | I483 | ICD10 | Inpatient | 34 |
| AF | I484 | ICD10 | Inpatient | 33 |
| AF | I489 | ICD10 | Inpatient | 5 |
| AF | 4273 | ICD9 | Inpatient | 65 |
| Other dementia | X0037 | Read code | GP | 3 |
| Other dementia | F111. | Read code | GP | 2 |
| Other dementia | Eu020 | Read code | GP | 1 |
| Other dementia | R043. | Read code | GP | 7 |
| Other dementia | X00Rk | Read code | GP | 2 |
| Other dementia | F11x0 | Read code | GP | 3 |
| Other dementia | F1440 | Read code | GP | 3 |
| Other dementia | F1420 | Read code | GP | 1 |
| Other dementia | F1422 | Read code | GP | 1 |
| Other dementia | F142z | Read code | GP | 2 |
| Other dementia | XE15P | Read code | GP | 33 |
| Other dementia | F11.. | Read code | GP | 1 |
| Other dementia | XE15F | Read code | GP | 4 |
| Other dementia | Xa0s2 | Read code | GP | 89 |
| Other dementia | A413. | Read code | GP | 1 |
| Other dementia | X0053 | Read code | GP | 6 |
| Other dementia | F281. | Read code | GP | 8 |
| Other dementia | F283. | Read code | GP | 17 |
| Other dementia | F29y3 | Read code | GP | 1 |
| Other dementia | XE0VM | Read code | GP | 8 |
| Other dementia | F1... | Read code | GP | 2 |
| Other dementia | F130. | Read code | GP | 3 |
| Other dementia | F1303 | Read code | GP | 3 |
| Other dementia | F1304 | Read code | GP | 29 |
| Other dementia | F1305 | Read code | GP | 1 |
| Other dementia | F130z | Read code | GP | 1 |
| Other dementia | F2... | Read code | GP | 4 |
| Other dementia | XaKyY | Read code | GP | 31 |
| Other dementia | X003A | Read code | GP | 11 |
| Other dementia | A411. | Read code | GP | 5 |
| Other dementia | XabVp | Read code | GP | 1 |
| Other dementia | F12.. | Read code | GP | 692 |
| Other dementia | F12z. | Read code | GP | 66 |
| Other dementia | 3314 | ICD9 | Inpatient | 5 |
| Other dementia | 3318 | ICD9 | Inpatient | 1 |
| Other dementia | F020 | ICD10 | Inpatient | 78 |
| Other dementia | F021 | ICD10 | Inpatient | 1 |
| Other dementia | F022 | ICD10 | Inpatient | 2 |
| Other dementia | F023 | ICD10 | Inpatient | 111 |
| Other dementia | F024 | ICD10 | Inpatient | 1 |
| Other dementia | F028 | ICD10 | Inpatient | 116 |
| Other dementia | F03 | ICD10 | Inpatient | 1180 |
| Other dementia | F09 | ICD10 | Inpatient | 15 |
| Other dementia | G310 | ICD10 | Inpatient | 99 |
| Other dementia | G311 | ICD10 | Inpatient | 3 |
| Other dementia | G312 | ICD10 | Inpatient | 103 |
| Other dementia | G318 | ICD10 | Inpatient | 290 |
| Other dementia | G319 | ICD10 | Inpatient | 641 |
| Hypercholesterolemia | C320. | Read code | GP | 123 |
| Hypercholesterolemia | XE11S | Read code | GP | 6545 |
| Hypercholesterolemia | C3200 | Read code | GP | 186 |
| Hypercholesterolemia | X40X0 | Read code | GP | 12 |
| Hypercholesterolemia | C320y | Read code | GP | 21 |
| Hypercholesterolemia | C320z | Read code | GP | 379 |
| Hypercholesterolemia | Xa9As | Read code | GP | 4808 |
| Hypercholesterolemia | 2720 | ICD9 | Inpatient | 27 |
| Hypercholesterolemia | 27209 | ICD9 | Inpatient | 2 |
| Hypercholesterolemia | E780 | ICD10 | Inpatient | 49849 |
| Hypertension | F282. | Read code | GP | 50 |
| Hypertension | G20.. | Read code | GP | 28 |
| Hypertension | XE0Uc | Read code | GP | 30795 |
| Hypertension | XM02V | Read code | GP | 2066 |
| Hypertension | XE0Ub | Read code | GP | 9795 |
| Hypertension | XaZWm | Read code | GP | 314 |
| Hypertension | Xab9L | Read code | GP | 143 |
| Hypertension | Xab9M | Read code | GP | 14 |
| Hypertension | XaZWn | Read code | GP | 1 |
| Hypertension | G2y.. | Read code | GP | 27 |
| Hypertension | G2z.. | Read code | GP | 761 |
| Hypertension | XaZbz | Read code | GP | 138 |
| Hypertension | 3482 | ICD9 | Inpatient | 1 |
| Hypertension | 4010 | ICD9 | Inpatient | 2 |
| Hypertension | 4011 | ICD9 | Inpatient | 1 |
| Hypertension | 4019 | ICD9 | Inpatient | 201 |
| Hypertension | G932 | ICD10 | Inpatient | 132 |
| Hypertension | I10 | ICD10 | Inpatient | 112951 |
| VaD | XE1Xs | Read code | GP | 139 |
| VaD | X003V | Read code | GP | 9 |
| VaD | Xa0lH | Read code | GP | 8 |
| VaD | X003T | Read code | GP | 4 |
| VaD | Eu01z | Read code | GP | 12 |
| VaD | Eu01y | Read code | GP | 2 |
| VaD | F019 | ICD10 | Inpatient | 478 |
| VaD | F011 | ICD10 | Inpatient | 23 |
| VaD | F013 | ICD10 | Inpatient | 6 |
| VaD | F010 | ICD10 | Inpatient | 5 |
| VaD | F018 | ICD10 | Inpatient | 5 |
| VaD | F012 | ICD10 | Inpatient | 2 |

**Suppl. Table 1**: Codes used to extract the diagnoses. The columns indicate (1) the disorder of interest, (2) the code corresponding to the disorder, (3) the coding system to which the code belongs, (4) whether the code was part of the inpatient- or the primary-care (GP) record, and (5) how many participants in UK Biobank were assigned a given diagnosis. All codes and coding systems were provided by UK Biobank, which also provides with tables for converting between the different coding systems. Note that any single patient might have been assigned several distinct codes (for the same disorder).

| **AF diagnosis** | **Ever prescribed warfarin** | |
| --- | --- | --- |
|  | **Yes (median age in years)** | **No (median age in years)** |
| **Yes** | 2,064 (64.9) | 1,386 (63.9) |
| **No** | 3,267 (63.4) | 97,904 (61.2) |

**Suppl. Table 2:** Frequencies and average ages at recruitment of participants diagnosed with AF and of participants with a history of warfarin use.

| **AF diagnosis** | **Ever prescribed warfarin** | |
| --- | --- | --- |
|  | **Yes (median age in years)** | **No (median age in years)** |
| **Yes** | 2,168 (64.7) | 1,536 (63.3) |
| **No** | 3,699 (62.3) | 133,419 (58.3) |

**Table 2:** Frequencies and average ages of participants diagnosed with AF and participants with a history of warfarin use.

**Suppl. Table 3:** Frequencies and average ages at recruitment of participants with different *VKORC1*-genotypes and participants diagnosed with general dementia, with ADem, and with VaD.

|  | **General dementia** | | |
| --- | --- | --- | --- |
|  |  | **Yes (median age in years)** | **No (median age in years)** |
| **Genotype (rs9923231)** | CC | 931 (65.3) | 81,781 (61.6) |
|  | CT | 1,119 (65.3) | 97,998 (61.5) |
|  | TT | 357 (65.3) | 29,237 (61.6) |
|  | **ADem** | | |
|  |  | **Yes (median age in years)** | **No (median age in years)** |
| **Genotype (rs9923231)** | CC | 323 (65.6) | 82,389 (61.6) |
|  | CT | 397 (65.4) | 98,720 (61.6) |
|  | TT | 124 (66.1) | 29,470 (61.6) |
|  | **VaD** | | |
|  |  | **Yes (median age in years)** | **No (median age in years)** |
| **Genotype (rs9923231)** | CC | 106 (67.0) | 82,606 (59.0) |
|  | CT | 135 (66.2) | 98,982 (59.0) |
|  | TT | 60 (66.9) | 29,534 (59.0) |

| **rs9923231** | **OR** | **95% CI** | **p** | **n cases** |
| --- | --- | --- | --- | --- |
| **Parental dementia** | | | | |
| CC | 1 (ref). |  |  | 12,178 |
| CT | 1.03 | 1.00-1.07 | 0.035 | 15,049 |
| TT | 1.08 | 1.04-1.13 | 2.3x10^-4^ | 4,654 |
| **General dementia** | | | | |
| CC | 1 (ref). |  |  | 931 |
| CT | 1.03 | 0.93-1.13 | 0.58 | 1,119 |
| TT | 1.09 | 0.96-1.25 | 0.19 | 357 |
| **ADem** | | | | |
| CC | 1 (ref.) |  |  | 323 |
| CT | 1.01 | 0.87-1.19 | 0.87 | 397 |
| TT | 1.13 | 0.91-1.40 | 0.27 | 124 |
| **VaD** | | | | |
| CC | 1 (ref.) |  |  | 106 |
| CT | 1.13 | 0.85-1.50 | 0.41 | 135 |
| TT | 1.72 | 1.20-2.44 | 0.0025 | 60 |

**Suppl. Table 4**: Logistic regression models with VaD as the outcome and rs9923231 as the predictor. Non-additive model with CC, CT, and TT as rs9923231 genotypes.

| **VaD** | | | | |
| --- | --- | --- | --- | --- |
| **Genotype** | **OR** | **95% CI** | **p** | **n cases** |
| Additive effect | 1.29 | 1.07-1.54 | 0.0060 | 301 |
| Hypertension | 3.21 | 2.38-4.36 | 4.8x10^-14^ | 224 |
| Hypercholesterolemia | 1.78 | 1.35-2.34 | 3.8x10^-5^ | 124 |

**Suppl. Table 5**: Logistic regression models with VaD as the outcome and rs9923231 as the predictor (as an additive effect), with the inclusion of hypertension and hypercholesterolemia as covariates.

| **Interaction** | **Covariate** | **Interaction effect** | | |
| --- | --- | --- | --- | --- |
|  |  | **Beta** | **SE** | **p** |
| AF*rs9923231 | warfarin use | 0.25 | 0.42 | 0.56 |
| rs9923231*warfarin use | AF | 0.0032 | 0.0050 | 0.77 |
| AF*warfarin use | *VKORC1* | -0.0059 | 0.0070 | 0.40 |

**Suppl. Table 6**: Models with VaD as the outcome and two-way interactions between rs9923231, AF, and warfarin use as predictors. The statistics represent the strength of the interaction between the first two variables with the inclusion of the third as a covariate.

| **VaD** | **Univariate** | | | **Multivariate** | | |
| --- | --- | --- | --- | --- | --- | --- |
| **Predictors** | **Beta_log-odds_** | **SE** | **p** | **Beta_log-odds_** | **SE** | **p** |
| rs9923231 | 0.25 | 0.091 | 0.0069 | 0.31 | 0.13 | 0.020 |
| warfarin use | 0.0029 | 0.0028 | 0.32 | -4.9x10^-4^ | 0.0037 | 0.90 |
| AF | 0.85 | 0.26 | 0.0011 | 0.87 | 0.33 | 0.0089 |

| **ADem** | **Univariate** | | | **Multivariate** | | |
| --- | --- | --- | --- | --- | --- | --- |
| **Predictors** | **Beta_log-odds_** | **SE** | **p** | **Beta_log-odds_** | **SE** | **p** |
| rs9923231 | 0.050 | 0.054 | 0.35 | 0.027 | 0.068 | 0.70 |
| warfarin use | -5.2x10^-5^ | 0.0021 | 0.98 | 5.5x10^-4^ | 0.0022 | 0.80 |
| AF | -0.053 | 0.22 | 0.81 | -0.16 | 0.24 | 0.49 |

**Suppl. Table 7**: Models predicting VaD (top), ADem (middle), and general dementia (bottom). For each outcome, the effects of three variables – rs9923231, warfarin use, and AF – are shown when each variable is predicting the outcome independently of the other two (univariate; left side), or in addition to the other two (multivariate, right side).

| **General dementia** | **Univariate** | | | **Multivariate** | | |
| --- | --- | --- | --- | --- | --- | --- |
| **Predictors** | **Beta_log-odds_** | **SE** | **p** | **Beta_log-odds_** | **SE** | **p** |
| rs9923231 | 0.041 | 0.032 | 0.21 | 0.024 | 0.041 | 0.57 |
| warfarin use | 0.0019 | 0.0010 | 0.069 | 5.8x10^-4^ | 0.0012 | 0.63 |
| AF | 0.52 | 0.10 | 6.1x10^-7^ | 0.35 | 0.12 | 0.0036 |

**Suppl. Figure 1:** Age distributions of different groups of participants at the time of recruitment to UK Biobank.

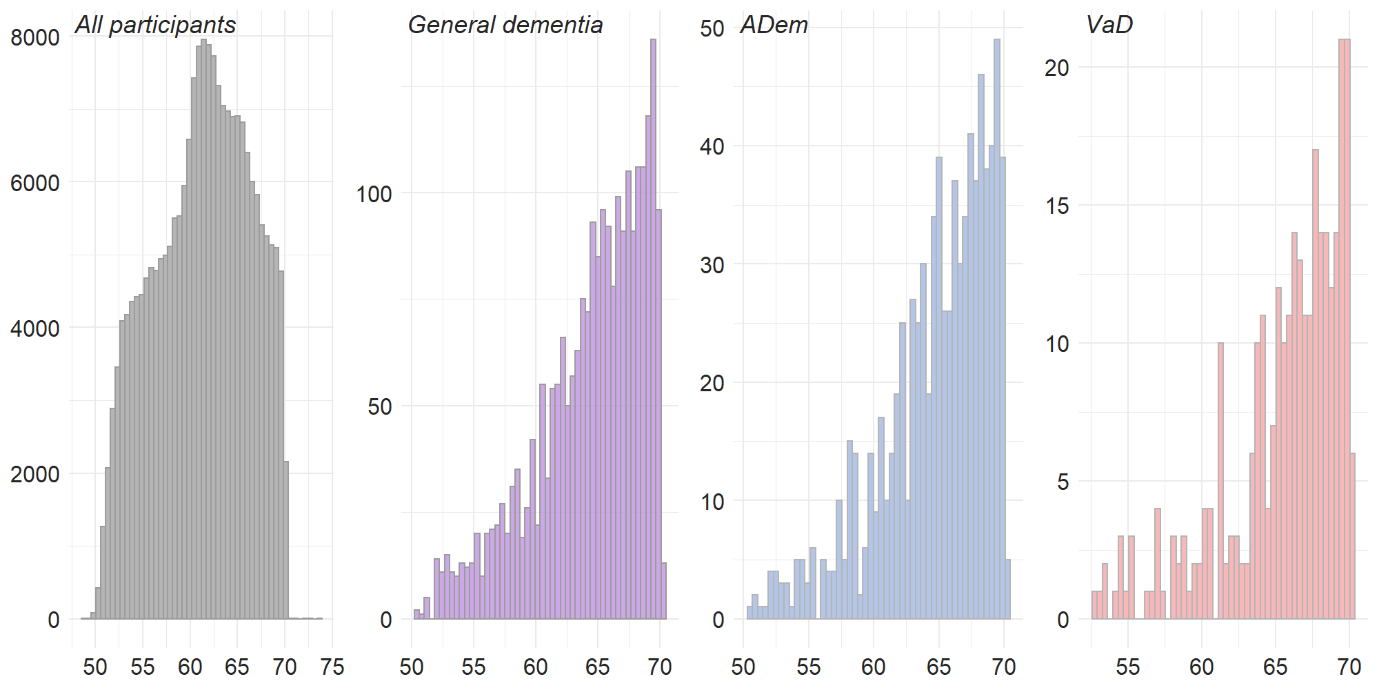

*n=844*

*n=211,423*

*n=301*

*n=2,407*

**Count**

**Age**

**Suppl. Text 1:** Power analysis for the interaction effect

A post hoc power analysis using as parameters an alpha error rate of 0.05, a sample size of 104,621, a case rate of VaD of 0.0014, and an effect allele frequency of 0.609 revealed that the power required to detect a two-way interaction effect (OR=1.1) between any two of the studied variables (warfarin use, AF, rs9923231) was approximately 5%. The power analysis was performed using the genpwr library in R.
